# scDeepC3: scRNA-seq Deep Clustering by A Skip AutoEncoder Network with Clustering Consistency

**DOI:** 10.1101/2022.06.05.494891

**Authors:** Gang Wu, Junjun Jiang, Xianming Liu

## Abstract

Single-cell RNA sequencing (scRNA-seq) reveals the heterogeneity and diversity among individual cells and allows researchers conduct cell-wise analysis. Clustering analysis is a fundamental step in analyzing scRNA-seq data which is needed in many downstream tasks. Recently, some deep clustering based methods exhibit very good performance by combining the AutoEncoder reconstruction-based pre-training and the fine-tune clustering. Their common idea is to cluster the samples by the learned features from the bottleneck layer of the pre-trained model. However, these reconstruction-based pre-training cannot guarantee that the learned features are beneficial to the clustering. To alleviate these issues, we propose an improved scRNA-seq Deep Clustering method by a skip AutoEncoder network with Clustering Consistency (i.e., named scDeepC3) from two aspects, an efficient network structure and a stable loss function. In particular, we introduce an adaptive shortcut connection layer to directly add the shallow-layer (encoder) features to deep-layer (decoder). This will increase the flow of forward information and back-forward gradients, and make the network training more stable. Considering the complementarity between the features of different layers, which can be seen as different views of the original samples, we introduce a clustering consistency loss to make the clustering results of different views consistent. Experimental results demonstrate that our proposed scDeepC3 achieves better performance than state-of-the-arts and the detailed ablation studies are conducted to help us understand how these parts make sense.

## 1 Introduction

Single-cell RNA sequencing (scRNA-seq) data contains numerous cells with thousands of gene expression, which can reveal the heterogeneity and diversity among individual cells and enable cell types identification analysis (1; 2; 3). In particular, it helps researchers conduct cell-wise analysis instead of bulk scRNA-seq data analysis, in which gene expression measurements are averaged over a population of cells. In order for researchers to make full use of these rich datasets, effective computational methods are needed. The computational analysis of scRNA-seq data involves multiple steps, including quality control, mapping, quantification, normalization, clustering, finding trajectories, and identifying differential expressed genes. Since clustering is a key step in defining cell types, in this paper we mainly focus on the clustering methods (4; 5; 6).

Clustering has been well studied in recent decades and many popular methods have been proposed, like K-Means (7), spectral clustering (8; 9; 10) and subspace clustering (11; 12). However, directly using these methods to cluster scRNA-seq data usually fails to obtain desirable results. This is mainly due the following statistical and computational challenges: (i) the low RNA capture rate leads to failure of detection of an expressed gene resulting in a dropout event, *i*.*e*., there are too many “false” zero count observation. (ii) scRNA-seq data usually contains highly dimensional features with thousands of genes so that common distance measurements are not effective because of the *Curse of Dimensionality* (13). (iii) compared to the bulk RNA-seq data and microarray data, scRNA-seq obtains count matrix on thousands of cells at once, which requires the clustering methods to be scale-able and efficient to handle large-scale data.

With the development of scRNA-seq analysis, some methods have been proposed to overcome these challenges. For example, Xu and Su proposed SNN-Cliq (14), which utilizes the concept of shared nearest-neighbor to effectively handle high-dimensional data. In (15), SC3 method introduces a pipeline for scRNA-seq clustering analysis. It integrates multiple clustering results through a consensus approach, and achieves very good performance. Wang et al. proposed a similarity-learning framework based on single-cell interpretation via multi-kernel learning (SIMLR) to obtain effective and stable distance metrics. Park et al. proposed a multi-kernel spectral clustering method (MPSSC) to impose the sparse structure on the similarity graph via *L*_1_ constraints (16). To obtain stable and meaningful distance metrics, dimensionality-reduction methods are usually leveraged to get composed features before clustering. t-distributed stochastic neighbor embedding (t-SNE) (17) is also widely used for reducing dimensions and embedding high-dimensional data for visualization. But it is not designed for scRNA-seq data and cannot address the issue of dropouts directly. Zero inflated factor analysis (ZIFA) method (18) is a dimensionality-reduction model that designed for scRNA-seq data. To address dropout counts, it implements a modified probabilistic principal component analysis based dimensionality-reduction method by incorporating a zero-inflated scheme. Then, clustering methods can be conducted on the low-dimension features to predict the final results.

Meanwhile, some other methods focus on imputing the false zeros caused by dropout in scRNA-seq data and try to improve the data quality for downstream analysis including traditional methods CIDR (clustering through imputation and dimensionality reduction) (19) and scImpute (20). In addition, DeepImpute (21) utilizes a deep neural network to predict dropout counts using highly correlated genes.

Most recently, deep learning has achieved great success in computer vision, nature language processing, and many other fields (22). Inspired by these work, many recent works in the field of scRNA-seq analysis resort to using artificial neural network to tackle the statistical and computational challenges (21; 23; 24; 25). DeepImpute (21) utilizes a standard fully connected deep neural network to estimate the dropouts in data. Deep count AutoEncoder (DCA) (23), based on AutoEncoder a widely used unsupervised model, replaces mean square error (MSE) loss with zero-inflated negative binomial (ZINB) model-based loss. DCA is effective at characterizing scRNA-seq and achieves better feature representation, which can benefit the downstream tasks. For clustering analysis, they usually extracted embedded features first and then conduct clustering method like K-Means to get final results separately. However, these two-stage approaches separate feature extraction and clustering, which cannot make sure the features extracted by the deep neural networks are cluster-friendly. Therefore, the clustering results are not the optimal. To this end, the single-cell model-based deep embedded clustering (scDeepCluster) (24) and deep embedding for single-cell clustering (DESC) (25) methods combine the feature extraction and clustering process and design an end-to-end clustering framework for scRNA-seq data. They train networks with the reconstruction loss and deep embedded clustering (DEC) loss (26) and get clustering results from the model directly.

Compared to traditional scRNA-seq clustering methods, these deep learning based methods usually achieve better performance. Because they have powerful representation and prediction ability and the fully connected structure is suitable for dimension-reduction and feature extraction naturally (27). In addition, it is also scalable for the large-scale dataset since the model optimization process uses standard back-propagation and stochastic gradient descent. The large-scale data can be easily processed by mini-batch training strategy which has been widely used. Finally, the joint reconstruction loss and clustering loss make the learned features more cluster-friendly, which further benefits downstream tasks (23; 24; 25; 26).

Usually, the deep clustering loss needs to be initialized with the clustering centroids. Considering that the deep clustering loss is conducted on the latent space (the bottleneck feature), to get meaningful initialization, pre-training strategy is used to transform the data into the feature space and then K-Means method is applied to these embedded features. Then clustering centroids can be seen as the initial solutions (weights) for deep clustering loss and the network can be trained with reconstruction and clustering losses jointly. Here, when the pre-trained model uses the ZINB model, it can be seen as the DCA method and then deep clustering loss is applied to fine-tune. However, in practice, we found that pre-training process has a great influence on the final clustering performance. As shown in Fig. 1, we can intuitively find that the clustering performance is unstable with different pre-trained model and sometimes the performance of deep clustering even has a negative effect. Because, on the one hand, initial features obtained in different pre-training latent space may be transition state or over-fitting, which reduce the effect of clustering loss. On the other hand, the reconstruction task in pre-training process and the clustering task are under different hypothesis, the reconstruction loss maps the data into a latent space where the feature is beneficial to reconstruct with as more information as possible, but this is not necessarily helpful to clustering task.

**Figure 1:**
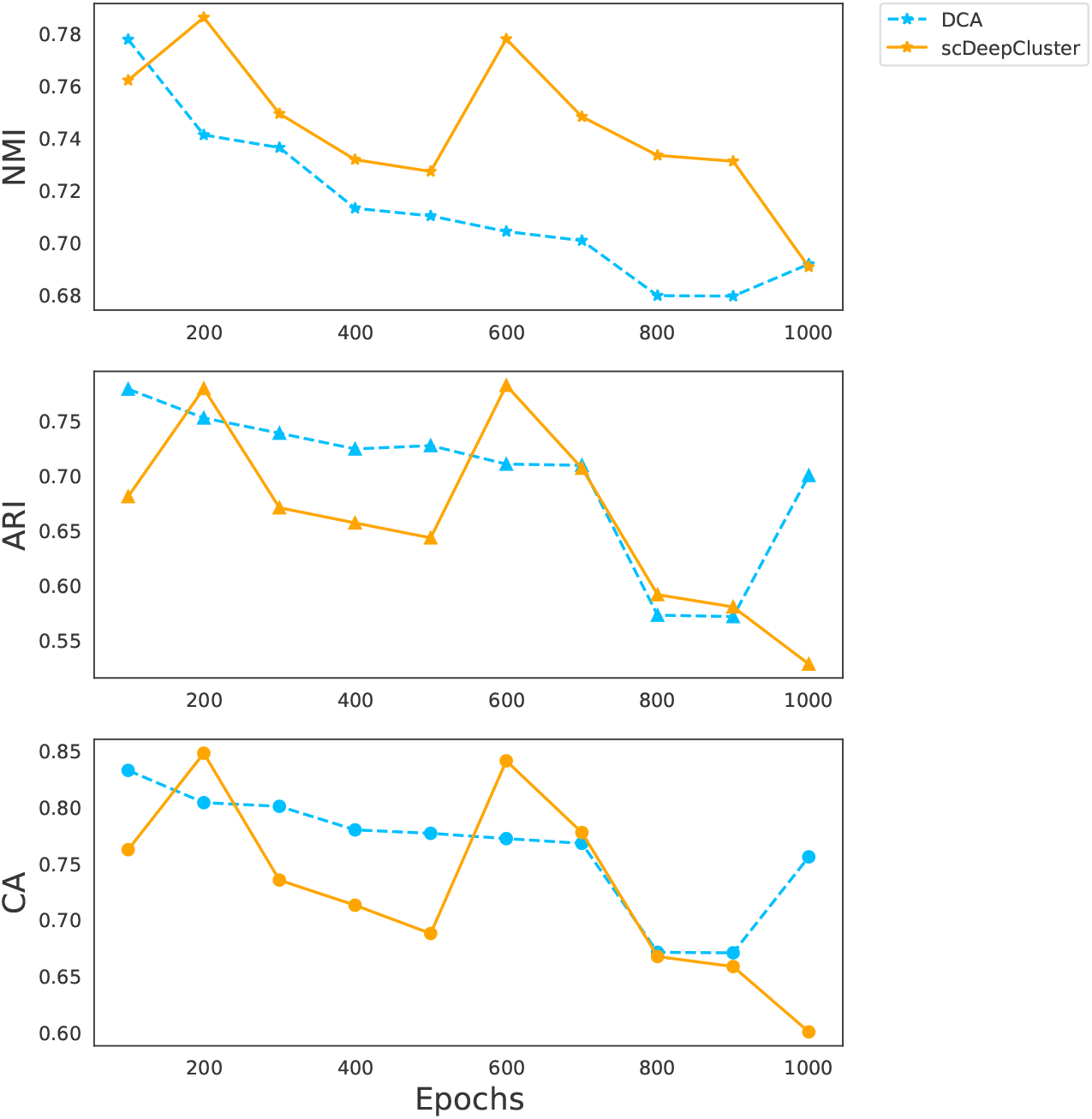
The clustering performance with different pre-training epochs. DCA represents the K-Means results of the pre-trained features. The scDeepCluster represents results after clustering fine-tune process.

In order to alleviate the above problems, in this paper we therefore introduce an adaptive shortcut connection layer and a deep clustering consistency loss to address these problems. Our framework is shown in Fig. 2. On the one hand, we improve the standard AutoEncoder with a simple, straight-forward but effective network structure to extract robust latent features. Here, we introduce an adaptive shortcut connection layer which adds the shallow-layer (encoder) features to deep-layer (decoder), so that we can expect to “see” the low-level feature representations during decoding (please refer to the red dash box in Fig. 2, and more details can be found at Section 3.2). This residual learning like structure can not only increase the flow of forward information and back-forward gradients, but also make the network training more stable and extract more general features. On the other hand, we think features extracted from different layers in fully connected neural network can be naturally regarded as multi-view features of the origin sample. Thus, the clustering loss can not only be conducted on the single bottleneck feature but also on the other features. They are all meaningful and the results of different views should be consistent. Therefore, we propose a multi-view clustering consistency constraint to simultaneously regularize the clustering results of the feature representations from the bottleneck layer and other layers in model. Considering the adaptive shortcut connection layer we introduced, we conduct the clustering consistency loss on the merged layer in the decoder to improve the single clustering loss with more information. This is reasonable especially when the pre-trained bottleneck features are not good (please refer to the blue dash box in Fig. 2, and more details can be found at Section 3.3).

**Figure 2:**
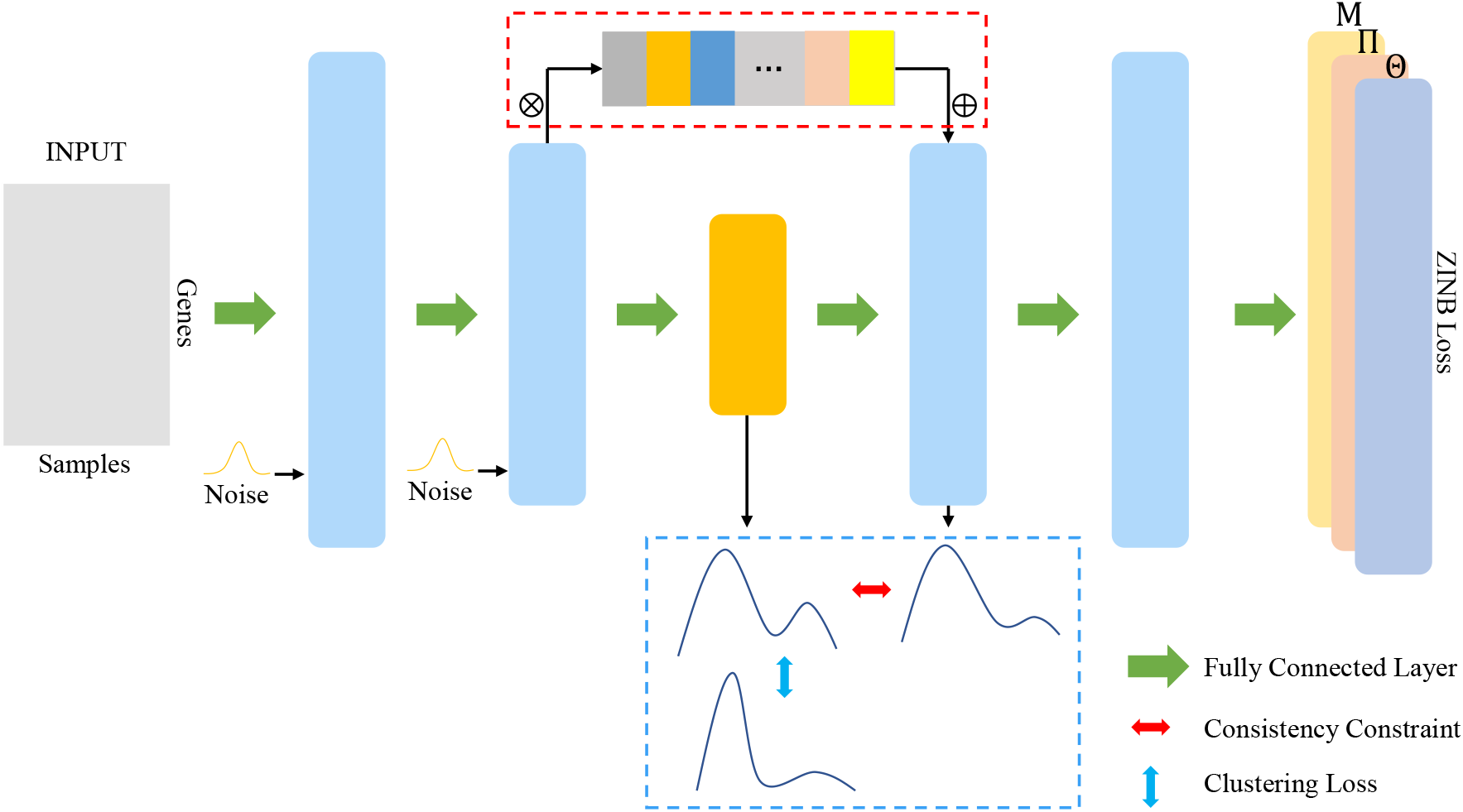
The overall network architecture of the proposed scDeepC3. “ × “ means element-wise multiplication and “+” means element-wise addition.

In summary, the contributions of this paper can be summarized as follows:

- We introduce a multi-view clustering consistency loss to improve the performance and stability of existing deep clustering loss. When compared with some existing methods, which conduct clustering with features from only the bottleneck layer, our method treats the features of different layers as different views of the sample and uses the multi-view features to regularize the clustering results.
- We improve the standard AutoEncoder model by adding an adaptive shortcut connection layer. On the one hand, this adaptive shortcut connection layer allows the important information in the input sample to be preserved. On the other hand, this residual learning like structure also increase the flow of forward information and back-forward gradients, and thus obtaining more stable results.
- Experimental results demonstrate that scDeepC3 achieves better performance when compared with the SOTAs. To intuitively understand how these parts make sense, we conducted detailed ablation studies.

The remaining parts of this paper are organized as follows: The Section 2 mainly reviews the representative works of scRNA-seq clustering. The Section 3 introduces the proposed scDeepC3 framework. The Section 4 contains the experiments, analysis and discussion of results. Finally, we summarize our work and look forward to the future work in the Section 5.

## 2 Related Work

In this section, we briefly review some methods that are most relevant to our work including *traditional methods* and *deep learning methods*.

### Traditional Methods

SIMLR (28) and MPSSC (16), based on spectral methods, mainly focus on the measure of the distance between samples and the multiple and learnable kernel functions are used to measure the distance between samples. Finally, stable and effective distance values are obtained. SIMLR and MPSSC are the most advanced measurement methods at this stage. These methods are spectral based which need to construct a full sample data matrix and a relationship matrix for processing. The calculation is complicated. Especially when processing large scale scRNA-seq data, MPSSC requires server-level hardware and it is unfriendly for individual researcher. SC3 (15) provides an easy-to-use algorithm tool library. It is based on spectral method and proposed a multi-view clustering boosting processing for scRNA-seq clustering analysis. SC3 achieves better and more stable performance through the boosting of the multi-view clustering results. In this paper, we combined the consistency of different clustering results under the multi-view feature of the same sample with the deep clustering loss. The consistency boosting process in SC3 is relatively simple which just use the voting method with the equal weights. Different from SC3, we combined the clustering consistency constraint with the deep embedding clustering loss in a simultaneously training process and achieved better performance. In general, the current traditional clustering methods based on spectral methods have achieved good results, but spectral methods have intensive calculation and high algorithm complexity. Due to the sharp increase in the number of scRNA-seq samples, how to efficiently handle large-scale Data is a big problem.

### Deep Learning Methods

Deep learning methods have made major breakthroughs in many fields and there have been many work applying deep learning model to scRNA-seq data clustering analysis task. DCA (23) took an AutoEncoder to extract low-dimension features. Contrast to the standard AutoEncoder, DCA combined the statistical information of the scRNA-seq data and designed the ZINB model, replacing the original MSE reconstruction loss, to model scRNA-seq data which effectively improved the extracted shallow features and achieved a better improvement with K-Means clustering methods. scDeepCluster (24) and DESC (25) are both end-to-end deep clustering methods which combined the feature extraction with the clustering using the deep clustering loss (26). Combining the feature extraction with the clustering loss makes the extracted features more cluster-friendly and further improves the clustering results. In addition, DESC demonstrates that the feature extraction under the guidance of the deep clustering model can make the features pay more attention to the cluster discrimination and effectively remove the batch effect. On the other hand, scVI (single-cell variational inference) (29) and scVAE (Single-cell variational AutoEncoders) (30) use the Variational AutoEncoder (VAE) (31) so that the model has the generation ability. In this paper, we focus on the deep clustering methods for clustering analysis of scRNA-seq data. Due to the ZINB loss has excellent effects in feature extraction, we are still based on the DCA framework, which is similar to scDeepCluster. Besides, we propose a deep clustering consistency loss and introduce an adaptive shortcut connection layer to further improve clustering performance.

## 3 Methodology

### 3.1 Overview

Recently, deep clustering methods have been extensively studied and been applied in scRNA-seq data, such as scDeepCluster (24) and DESC (25), building an end-to-end deep clustering model to remove batch effect and cluster analysis tasks. Although the deep clustering strategy achieves better results than the traditional two-phase methods which have dimensionality reduction followed by clustering task, the deep clustering process is unstable due to the initialization of the weights of clustering loss requires K-Means method to obtain the clustering centroids. In addition, the existing deep learning methods such as scDeepCluster, DESC, scVI, and scVAE, usually cannot include very deep network structure, because too deep model is easily to overfit.

In order to solve the above two problems, we design a novel deep learning model for scRNA-seq data clustering analysis: (i) We add the multi-view feature clustering consistency constraint to the clustering loss to further improve the stability and accuracy of the clustering process (see Section 3.3). (ii) We design a new AutoEncoder structure, which introduces an adaptive shortcut connection layer to further improve the extracted features (see Section 3.2).

### 3.2 Network Architecture

#### ZINB Based Denoising AutoEncoder

The AutoEncoder is a widely used deep neural network for the unsupervised representation learning task (32) which was shown to outperform traditional approaches as it can learn complex structure in the data. An AutoEncoder usually contains three components: an encoder, a bottleneck layer and a decoder. The encoder is a stack of some fully connected layers activated by activation function and the decoder is reverse of encoder. The AutoEncoder learns in an unsupervised way to efficiently compress and subs reconstruct the data with typical MSE loss function. It is characterized by the fact that both input and output layers are of the same size and the bottleneck layer has much lower dimensions. In addition, denoising AutoEncoder (33) further improves the generality of the AutoEncoder. In contrast to AutoEncoder, the denoising AutoEncoder receives corrupted data as input and is trained to reconstruct the original uncorrupted data. Denoising AutoEncoder is proved to be more powerful to learn a robust compressed representation of data due to its ability of learning the representations of the input that are corrupted by small irrelevant changes in input. Here, we use the denoising AutoEncoder as our framework to map the input of read counts to an embedded space, as shown in Fig. 2. In practice, we use random Gaussian noise and add it into input data. Therefore, the input of the AutoEncoder is:

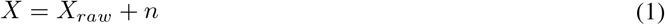

where *n* is the random Gaussian noise. Note that the noise is incorporated into every layer of the encoder, which is the same with the stack denoising AutoEncoder. Here, we define the nonlinear mapping of encoder as 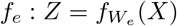 and the mapping of decoder is 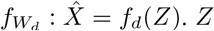 is the embedded features and *W*_*e*_ and *W*_*d*_ are the weights in the encoder and decoder. The learning process of the AutoEncoder minimizes the loss function

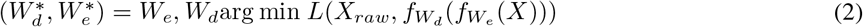

where *L* is the loss function.

Because the variance of gene expression in samples is often larger than its corresponding mean and the frequent dropout events (resulting from RNA capture inefficiency) will cause the count matrix containing amounts of zero values. For scRNA-seq data, in the paper, we apply the ZINB loss (23) to train the denoising AutoEncoder instead of MSE loss, as shown in Fig. 2. The loss function of ZINB is the likehood of a ZINB distribution, which can model highly sparse and overdispersed count data (23). Here, ZINB distribution is parameterized with mean (*μ*) and dispersion (*θ*) parameters of the negative binomial component and the mixture coefficient (*π*) that represents the probability of dropout events. In general, we use the deep neural networks to estimate the three parameters of ZINB distribution by the training data

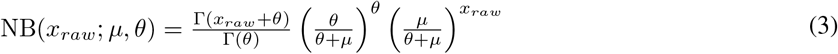

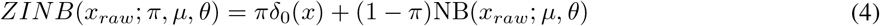

where *x*_*raw*_ is the original data without Gaussian noise. The denoising AutoEncoder estimates the mean (*μ*), dispersion (*θ*) and the drop coefficient (*π*) parameters. Note the last hidden layer of decoder as *D* and three fully connected layers are followed to estimate the parameters

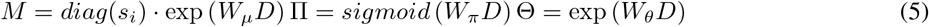

where *s*_*i*_ is size factors calculated from the input and *M*, Θ and Π represent the matrix form of estimations of mean, dispersion and dropout probability, respectively. Considering the coefficient Π should be limited to the value between zero and one, we use the sigmoid function as activation. The outputs of the mean and dispersion are exponential because the mean and dispersion parameters are always non-negative values. Therefore, the loss function uses the negative log of the ZINB likelihood:

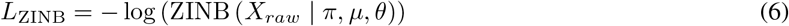

#### Adaptive Shortcut Connection Module

Inspired by the ResNet (34) and U-Net (35), the shortcut connection plays an important role in both works which makes training processing more stable and helps neural network to extract better features. In addition, shortcut connection fuses the low-level information in shallow layers and high-level semantic information in deep layers and this makes it possible to directly interact with features at different layers.

Based on the above observation, for the scRNA-seq clustering analysis, we design the adaptive shortcut connection layer and extend standard AutoEncoder structure with shortcut connection layer. Details of the adaptive shortcut connection layer is shown Fig. 3. It should be noted that the last layer in encoder is *H*^*e*^, the bottleneck feature is *Z* and the first layer in decoder is *H*^*d*^. The adaptive shortcut connection can be mathematically formulated as followings

**Figure 3:**
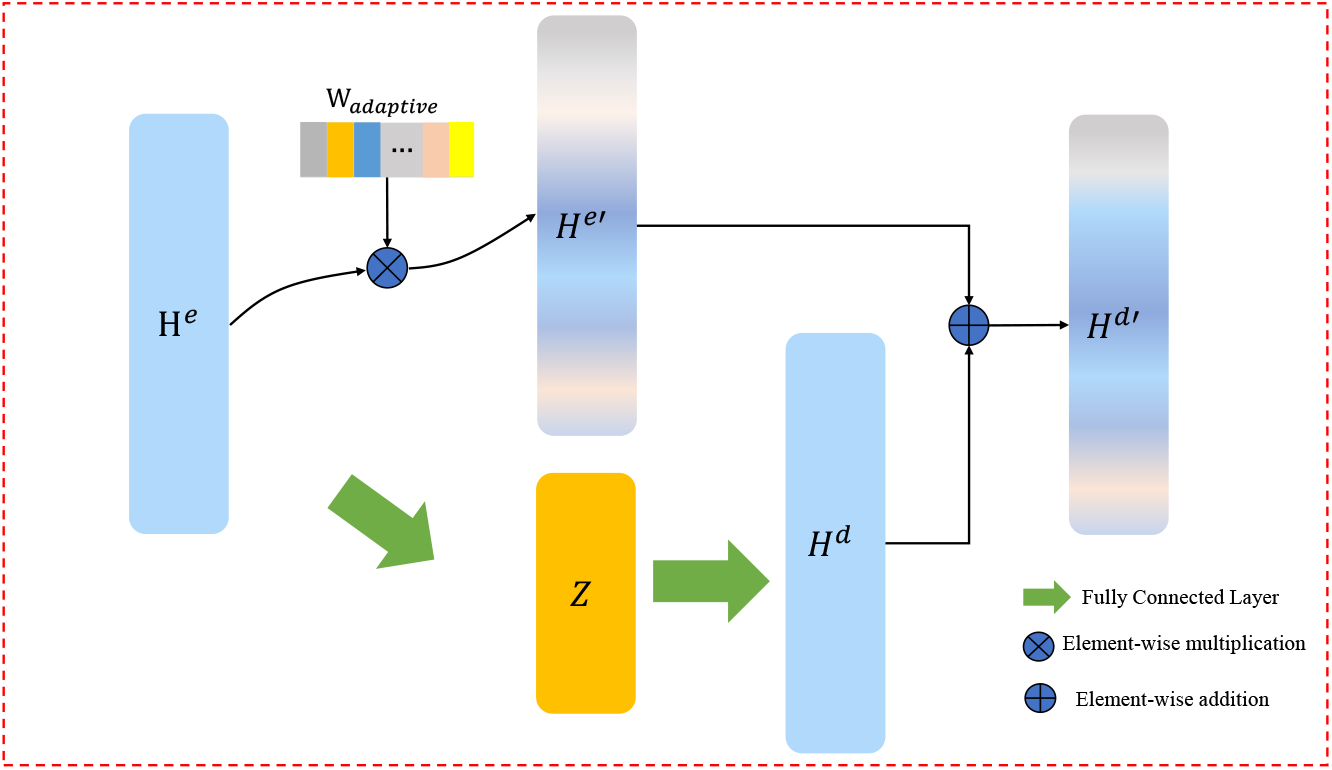
The detailed view of adaptive shortcut connection module.

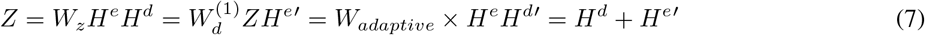

where *W*_*Z*_ is the weight of fully connected layers for latent feature and 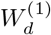 is the weight of the first fully connected layer in decoder. *W*_*adaptive*_ is the weight vector in the shortcut connection layer which is learnable and updated by neural networks and *×* means the element-wise multiplication. The obtained 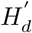 contains the original features and the information from output features of decoder and is fed into the following layers in the decoder. Through the skip connection, which fuses the shallow features of the encoder and the high-level features, it makes the current layer have richer information which helps the realization of clustering consistency loss.

### 3.3 Clustering Consistency Loss Function

#### Deep Embedded Clustering Loss

For the deep clustering stage, we apply the DEC loss (26) which is also used in (24) and (25). Consider the problem of clustering the set of *n* cells *X* with each sample *x*_*i*_ ∈ N^*d*^ (*x*_*i*_ represents the read counts of *d* genes in the *i*th cell) into *k* clusters. Instead of clustering directly in data space *X*, deep embedded clustering first finds a nonlinear mapping *f*_*e*_ : *x*_*i*_ → *z*_*i*_ where *Z* is the latent feature space and the dimension of *Z* is typically much smaller than *X* in order to avoid “curse of dimensionality” (13). Then the clustering process will be conducted on the latent space *Z*.

To capture the characters of scRNA-seq data, we use the denoising ZINB model to learn the nonlinear mapping from *X* to *Z* and apply DEC loss on the latent space *Z*. The cluster algorithm is the same as in (26), which is defined as Kullback–Leibler (KL) divergence between the distributions *P* and *Q*, where *Q* is the distribution of soft assignments calculated by Student’s t-distribution and *P* is the derivation of the target distribution from *Q*. Formally, the clustering loss is:

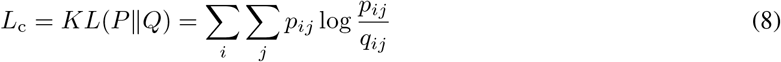

where *q*_*ij*_ is the soft label of embedded feature *z*_*i*_ which can be seen as the similarity between *z*_*i*_ and cluster centre *c*_*j*_ measured by Student’s t-distribution:

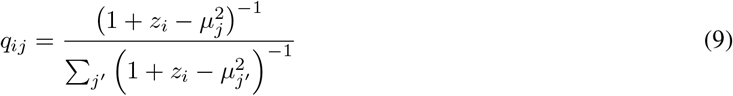

the target distribution *p*_*ij*_ is calculated with *q*_*ij*_:

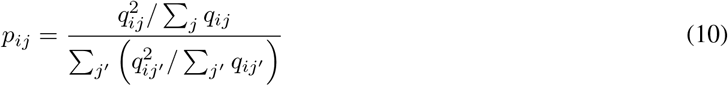

The training process is self-supervised because the target *P* is defined based on *Q*. The initialization of cluster centers needed in clustering loss is obtained by standard K-Means clustering. To get the meaningful clustering results we need to pre-train the ZINB model and then use the clustering loss to fine-tune.

#### Consistency Constraint

The deep embedding clustering loss is widely used in many clustering analysis works. However, in practice, we find that the training process is unstable and it is hard for us to choose a desirable model especially when we apply it to some new datasets. Because the training process for the deep clustering model is unsupervised and we have no label information for evaluation. Thus, the capacity and structure of the neural network directly determine how the features are extracted. The clustering loss is conducted on the latent feature space after pre-training. Therefore, the initialization of clustering loss will directly affect the fine-tune process and get different clustering result. Usually, the K-Means clustering method is conducted on latent space to initialize cluster centroids. However, we find with different initialization the results will be different and unstable.

To this end, in this paper, we propose a clustering consistency loss. After pre-training, the existing clustering loss is only conducted on the bottleneck layer. However, features in other layers can be seen as the different views for the same sample and the clustering results with these features should be consist. Here, we apply the KL divergence to measure the clustering consistency loss to make soft assignments (*Q*) be similar as *Q*′ calculated from other views. As shown in Fig. 4. We denote the features extracted from other layers as *H* and the soft assignment is measured by

**Figure 4:**
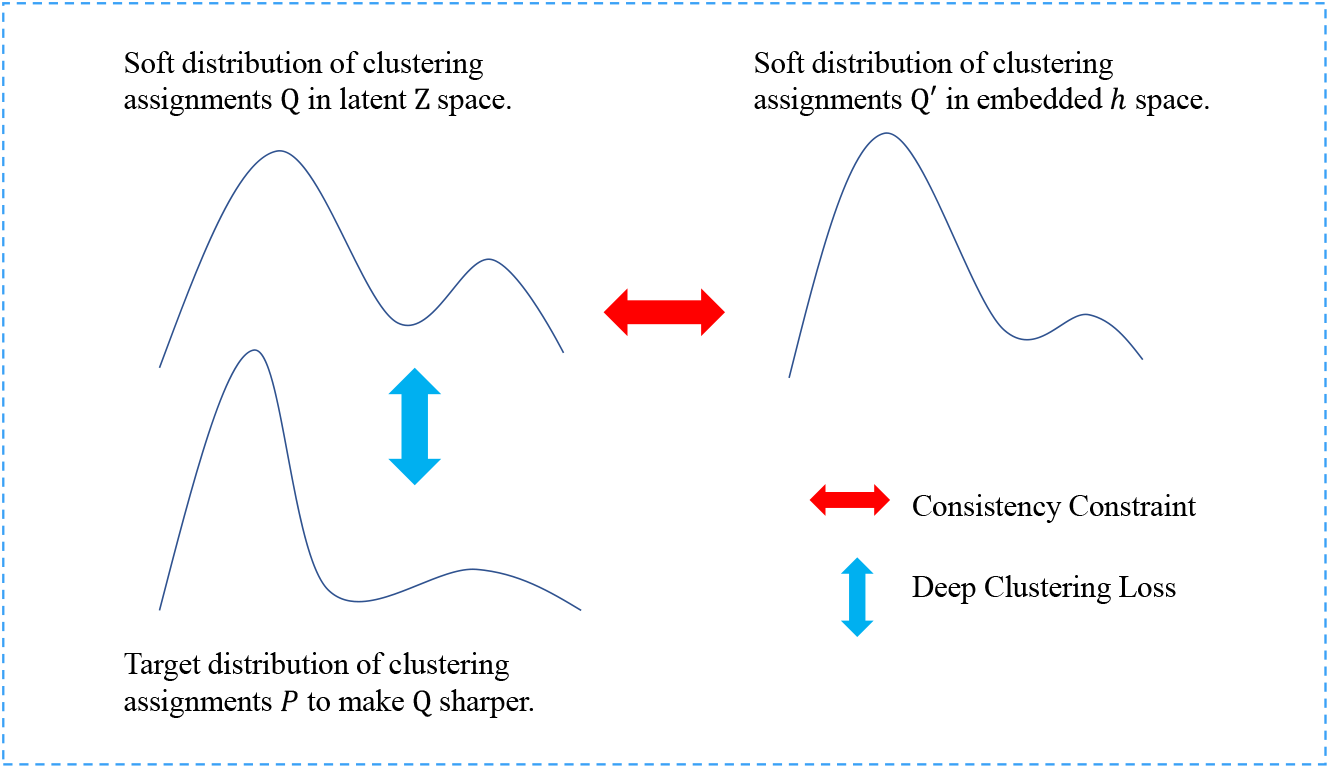
The detail view of clustering with consistency constraint module.

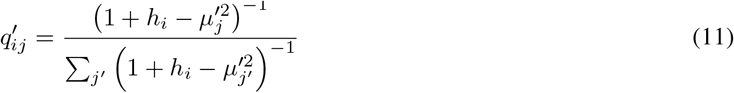

where *h*_*i*_ is the output feature of the other layer and 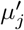 is the *j*-th cluster centroids in *H* feature space. We think the *q*_*ij*_ and 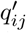 for the same sample should be consistent and we also use the KL divergence to define the clustering consistency constraint loss

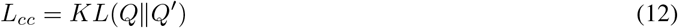

In practice, we choose the first layer in decoder as the consistency feature which is the output of adaptive shortcut connection and has more information about the sample. Note that the soft label *Q*′ used in KL divergence is aligned with *Q* using the Hungarian algorithm (36). Because in different feature space the order of cluster centroids will be different.

Finally, our model consists two components: the denoising ZINB model with adaptive shortcut connection layer and the deep clustering loss with the consistency constraint, shown in Fig. 3 and Fig. 4, respectively. The total objective function for the clustering is

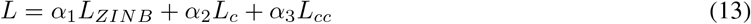

where *L*_*ZINB*_, *L*_*c*_ and *L*_*cc*_ are the ZINB loss, deep clustering loss and clustering consistency loss, respectively. *α*_1_, *α*_2_, *α*_2_ are the coefficients that control the relative weights among these losses.

### 3.4 Implementation Details

#### Data Pre-processing

The datasets used in this paper are the same with (24). For neural network numerical optimization stability, we need to transform discrete data into continuous smooth data. In particular, we first normalize the data matrix and then we take a natural log transformation on data. Finally, we transform the logarithm data into z-score data, which implies that each gene has zero mean and unit variance.

#### Hyper-Parameter Settings

We implement in Python3 using Keras with Tensorflow (37) backend. ADAM optimizer (38) is applied with an initial learning of 1e-4. In our experiments, we firstly pre-train the AutoEncoder with adaptive shortcut connection layer using the ZINB Loss. In contrast to many previous work which usually pre-train the network for 400, 600 or more epochs, we find our model will take fewer epochs to achieve a stable and desirable performance. For the fine-tune stage, we train the model with ZINB Loss, clustering loss and consistency constraint loss and the coefficients *α*_1_, *α*_2_ and *α*_3_ are 1.2, 1.2 and 1, respectively. We stop the training process when the percent of delta samples is less than 0.003. The batch size is 256. The size of fully connected hidden layers in the encoder are set to 256 and 64 and the decoder is the reverse of the encoder, e.g., 64 and 256. The bottleneck layer has a size of 32 and the adaptive shortcut connection layer has the same size of the last encoder layer which is 64 in our experiment.

## 4 Experiments and Results

In this section, we test our scDeepC3 on four real world scRNA-seq datasets. To verify its superiority, we compare the clustering performance with some conventional scRNA-seq data clustering algorithms, including SIMLR (28), SC3 (15) and some deep learning methods nameed DCA (23), DESC (25), scDeepCluster (24) and scVI (29). Other methods, like scVAE (30), are not in consideration of comparison in this section. Because we think scVAE and scVI have the similar framework for clustering analysis. They are both generative model based on VAE and combines the ZINB loss to extract latent features and then conduct the clustering methods. We use the cell type information provided by authors as ground truth to measure the accuracy of different methods. Then, we apply our method to get a clustering result and compare its similarity with the true label information. The comparison of these methods are summarized in section 4.1. Moreover, to better understand how scDeepC3 works with the adaptive shortcut connection layer and the clustering loss with consistency constraint, we conduct the ablation studies in 4.2 which demonstrate that both components we designed are effective.

### Implementation

The proposed method is available at GITHUB (we will release the code upon the acceptance of this paper). The other compared methods are tested with the official implementation to conduct clustering analysis.

### Evaluation Measures

All clustering results are measured by NMI (39), ARI (40) and CA.

The first one is normalized mutual information (NMI) (39):

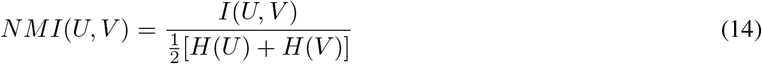

where *U* and *V* are two clustering assignments. *I* is the mutual information metric and *H* is entropy.

The second one is the adjusted rand index (ARI). It is a simple measure of agreement between two cluster assignments U and V. The ARI is calculated using four quantities. Specifically, we define:

- a: the number of pairs of two objects in the same group in both U and V
- b: the number of pairs of two objects in different groups in both U and V
- c: the number of pairs of two objects in the same group in U but in different groups in V
- d: the number of pairs of two objects in different groups in U but in the same group in V.

Base on the above quantities, the ARI is defined as:

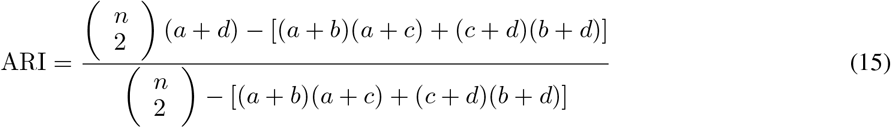

The last one is unsupervised clustering accuracy (CA):

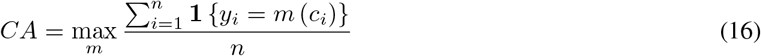

where *y*_*i*_ is the ground-truth label, *c*_*i*_ is the cluster assignment generated by the algorithm, and *m* is a mapping function which ranges over all possible one-to-one mappings between assignments and labels. It is obvious that this metric finds the best matching between cluster assignments from a clustering method and the ground truth. The optimal mapping function can be efficiently computed by the Hungarian algorithm (36).

### 4.1 Analysis of Real Data

The comparison of our approach and other competing methods are conducted on the several real world scRNA-seq datasets. We use the four real datasets introduced in (24) as benchmark. The four datasets were generated from four representative sequencing platforms: PBMC 4k cells from the 10X genomics platform (10X PBMC), mouse embryonic stem cells from a droplet barcoding platform (mouse ES cells), mouse bladder cells from the Microwell-seq platform (mouse bladder cells) and worm neuron cells from the scRNA-seq platform (worm neuron cells). The four datasets, respectively, had 4,271, 2,717, 2,746 and 4,186 cells per sample, with 16,449, 24,046, 19,079 and 11,955 genes after pre-processing, and form 8, 4, 16 and 10 groups per cluster, as summarized in Table 1. The experiments discussed mainly compares the four benchmark datasets to demonstrate the effectiveness of our proposed clustering loss with consistency constraint. Results are evaluated by NMI, ARI and CA.

**Table 1:** Summary of four real scRNA-seq datasets

| Dataset | Sample size/cell numbers | No. of genes | No. of groups |
| --- | --- | --- | --- |
| 10X PBMC | 4271 | 16449 | 8 |
| Worm neuron cells | 4186 | 11955 | 10 |
| Mouse bladder cells | 2746 | 19079 | 16 |
| Mouse ES cells | 2717 | 24046 | 4 |

Firstly, Fig. 5 shows the overall performance of 6 methods in 4 real datasets. We can see that our method has a better performance in all datasets. We can intuitively see that our method has a better performance than scDeepCluster and has the best results on 3 datasets. For compared traditional statistical-based methods, SIMLR provides relatively competitive performance. However, it performs poorly on some specific datasets such as Mouse ES data which just has 4 classes but has over 20 thousand gene features and has the NMI and ARI value *<*0.2.

**Figure 5:**
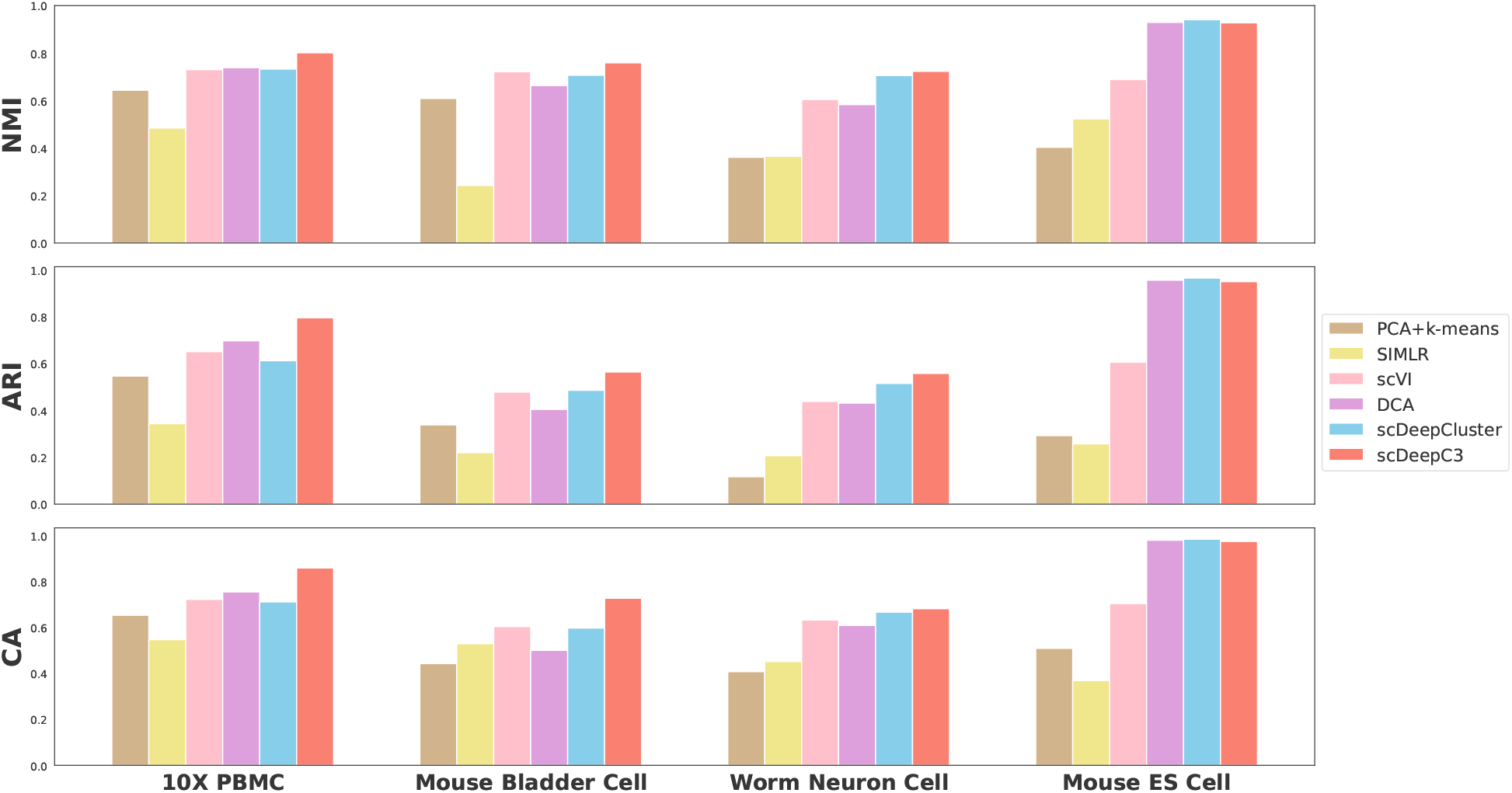
Benchmark results on four scRNA-seq datasets with true labels. Comparison of clustering performances of SIMLR (28), DCA (23), scVI (29), scDeepCluster (24), DESC (25), and ours, measured by NMI, ARI and CA.

To intuitively compare different clustering results, we use dimension reduction method (such as t-SNE (17)) to get 2D features. For our method, we extract the embedded features of the input data and apply t-SNE method with the default parameters to get 2D features. 2D features embedded by DCA and scDeepCluster are obtained in the same way. As shown in Fig. 6, we can find that DCA method can’t separate features clearly and our method scDeepC3 have the best performance. Especially, for 10X PBMC data, scDeepCluster can’t separate the yellow, green and brown samples and scDeepC3 separates them directly. For the worm neuron cell data, scDeepCluster separates overmuch and divides green samples into several sub-clusters. scDeepC3 obtains complete green samples with one cluster. Therefore, from an intuitive point of view, our method achieves better performance than DCA and scDeepCluster methods in these real world datasets.

**Figure 6:**
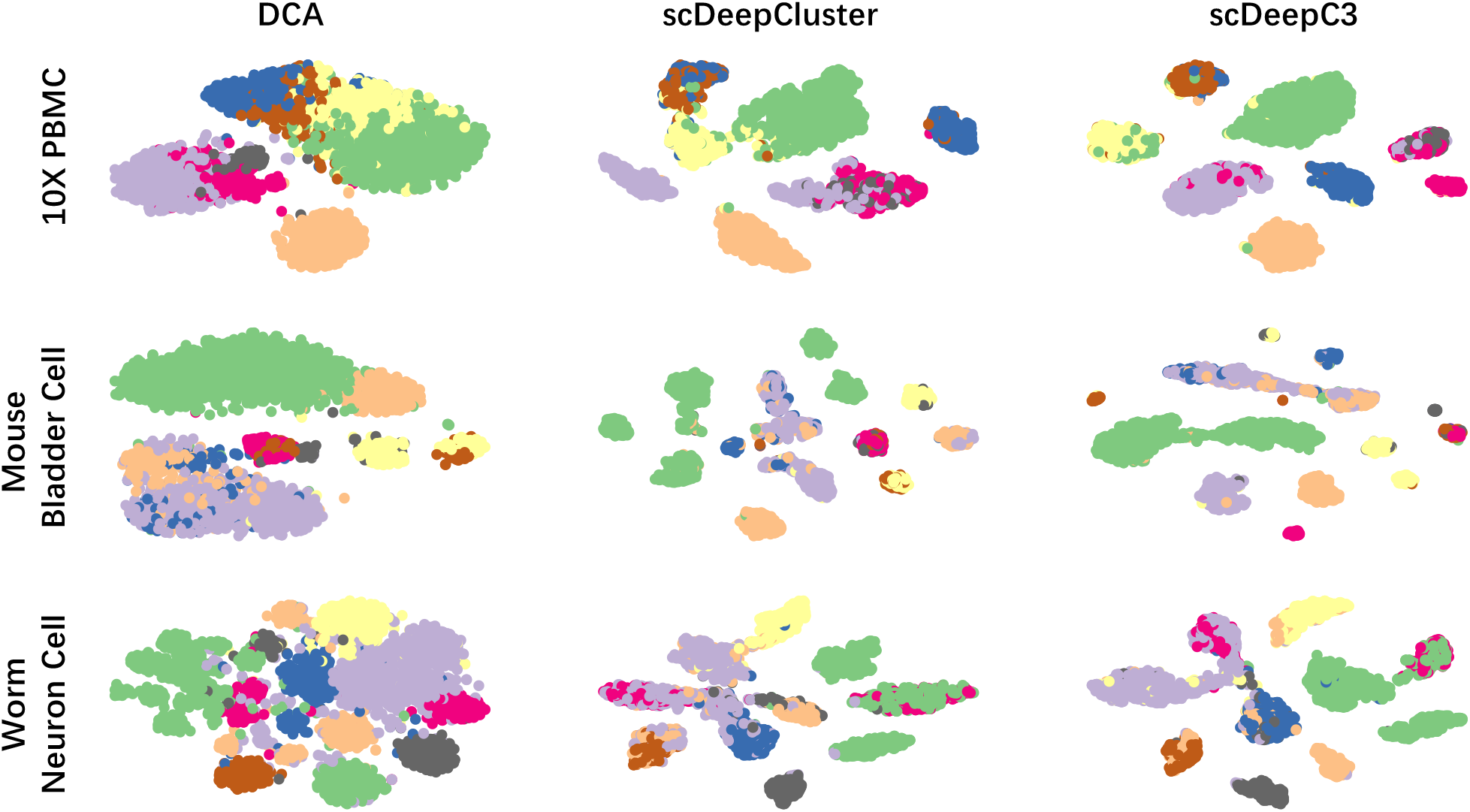
Visualization of embedded representations extracted by DCA (23), scDeepCluster (24) and our method scDeepC3. Each point represents a cell and is colored with distinct colors with label information.

### 4.2 Ablation Studies

The two main contributions of this paper are (i) designing a more effective AutoEncoder structure with adaptive shortcut connection layer and (ii) introducing a novel clustering loss to make the clustering results of different layer to be consistent. In this section, we conduct ablation experiments to verify these points.

#### Effectiveness of Adaptive Shortcut Connection Layer

Firstly, to demonstrate the effect of adaptive shortcut connection layer, we evaluate the clustering results of the features extracted by different AutoEncoder structures with or without the shortcut connection layer. In experiment, the standard AutoEncoder without shortcut connection structure is as the benchmark. To avoid intuitively reflect the how stable the pre-training process is, we use K-Means method to get clustering results directly conducted on features extracted from both model replacing the following clustering loss fine-tuning process. We train the model with 1000 epochs by 10 times and the mean results of these two model with different structure on four datasets are shown in Table. 2. We can find that, with the adaptive shortcut connection layer, we can achieve better results. Especially, for the Mouse bladder cell data, our method obtains almost 10 presents improvement on the clustering accuracy. To demonstrate this structure can improve the stability and generality of the model, as shown in Fig. 7, we report the intermediate results (are record every 100 epochs) of the training process on 10X PBMC, Mouse bladder bell and Worm neuron cell datasets. From these results, we can find that features extracted by our model have better and more stable performance. Especially for 10X PBMC data, the CA values reported in our model stable during this train process. In contrast, the model without shortcut connection layer will be sharply overfitting after hundreds of epochs. Due to the unsupervised training process, model having a robust and general training process is important and necessary.

**Table 2:** Benchmark on four dataset with or without the adaptive shortcut connection layer. And “w/o adaptive shortcut connection” is the basic AutoEncoder structure. “w. adaptive shortcut connection” is the framework our used in this work.

| Dataset | w/o adaptive shortcut connetion |  |  | w. adaptive shortcut connetion |  |  |
| --- | --- | --- | --- | --- | --- | --- |
|  | NMI | ARI | CA | NMI | ARI | CA |
| 10X_PBMC | 0.7456 | 0.7074 | 0.7653 | 0.7643 | 0.7785 | 0.8264 |
| mouse_bladder_cell | 0.6706 | 0.4096 | 0.5074 | 0.7163 | 0.4815 | 0.6046 |
| worm_neuron_cell | 0.5963 | 0.4431 | 0.6177 | 0.6750 | 0.4924 | 0.6304 |
| mouse_ES_cell | 0.9182 | 0.9378 | 0.9645 | 0.8920 | 0.8911 | 0.9231 |
| Mean | 0.7327 | 0.6245 | 0.7137 | 0.7619 | 0.6609 | 0.7461 |

**Figure 7:**
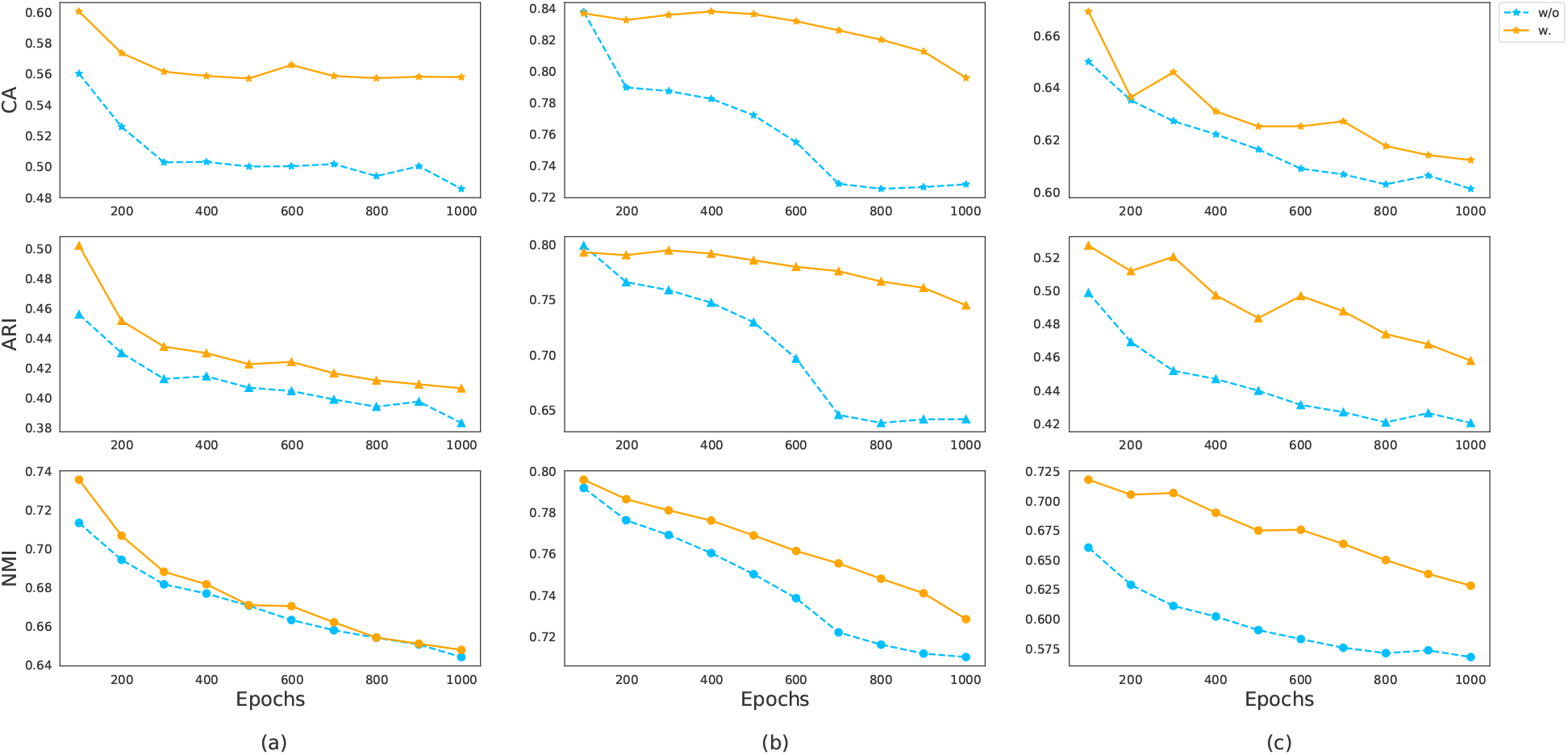
Comparison of clustering performances of different AutoEncoders with or without the adaptive shortcut connection. The results of the three datasets named Mouse bladder cell, 10X PBMC and Worm neuron cell are shown in (a), (b) and (c), respectively.

In summary, the proposed adaptive shortcut connection layer can extract more clustering-friendly features and has greater stability in training process.

#### Effectiveness of Clustering Loss with Consistency Constraint

In this subsection, we compare the fine-tuned process with or without the consistency constraint. The benchmark deep clustering loss we used is DEC loss (26), which is used in DESC (25) and scDeepCluster (24). To avoid being affected by different network structures, we first pre-trained the standard AutoEncoder model with ZINB loss and then used deep clustering loss with or without our consistency constraint to fine-tune the model. The results of both methods on 4 real datasets are shown in Table 3. We can find that our method with consistency loss achieved better performance on 3 real world datasets.

**Table 3:** Benchmark on four dataset with or without consistency constraint. K-Means represents the results of the K-Means method conducted directly on the extracted features. And “w/o consistency constraint loss” can be seen as the scDeepCluster using the deep clustering loss on single latent feature space. “w.consistency constraint loss” is our proposed loss.

| Dataset | K-Means |  |  | w/o consistency constraint loss |  |  | w. consistency constraint loss |  |  |
| --- | --- | --- | --- | --- | --- | --- | --- | --- | --- |
|  | NMI | ARI | CA | NMI | ARI | CA | NMI | ARI | CA |
| 10X_PBMC | 0.7456 | 0.7074 | 0.7653 | 0.7474 | 0.6501 | 0.7417 | 0.7620 | 0.7080 | 0.7796 |
| mouse_bladder_cell | 0.6706 | 0.4096 | 0.5074 | 0.7067 | 0.4782 | 0.5890 | 0.7104 | 0.4791 | 0.5931 |
| worm_neuron_cell | 0.5963 | 0.4431 | 0.6177 | 0.7121 | 0.5276 | 0.6750 | 0.7080 | 0.5124 | 0.6683 |
| mouse_ES_cell | 0.9182 | 0.9378 | 0.9645 | 0.9279 | 0.9455 | 0.9682 | 0.9301 | 0.9467 | 0.9691 |
| Mean | 0.7327 | 0.6245 | 0.7137 | 0.7735 | 0.6503 | 0.7435 | 0.7776 | 0.6616 | 0.7525 |

Considering clustering analysis is an unsupervised task and usually we cannot know in advance whether learning is enough without the ground truth. Therefore, a good clustering algorithm should be able to obtain stable results, and the results should not change with time (iteration times). In order to verify the proposed method intuitively, we report the results of our method (with different losses) at different training epochs. Especially, the comparison results of the 10X PBMC data are shown in Fig. 8. We can find that the embedded features cannot be assigned well by DEC loss, the clustering consistency constraint has better results, i.e., an increase of two percentage points. The results of our proposed clustering consistency loss are more stable during training process while scDeepCluster is unstable especially from epoch 200 to 400. This indicates that the loss with consistency constraint can obtain much more information from different feature spaces which can help clustering processing especially when the extracted features are unstable or overfitting.

**Figure 8:**
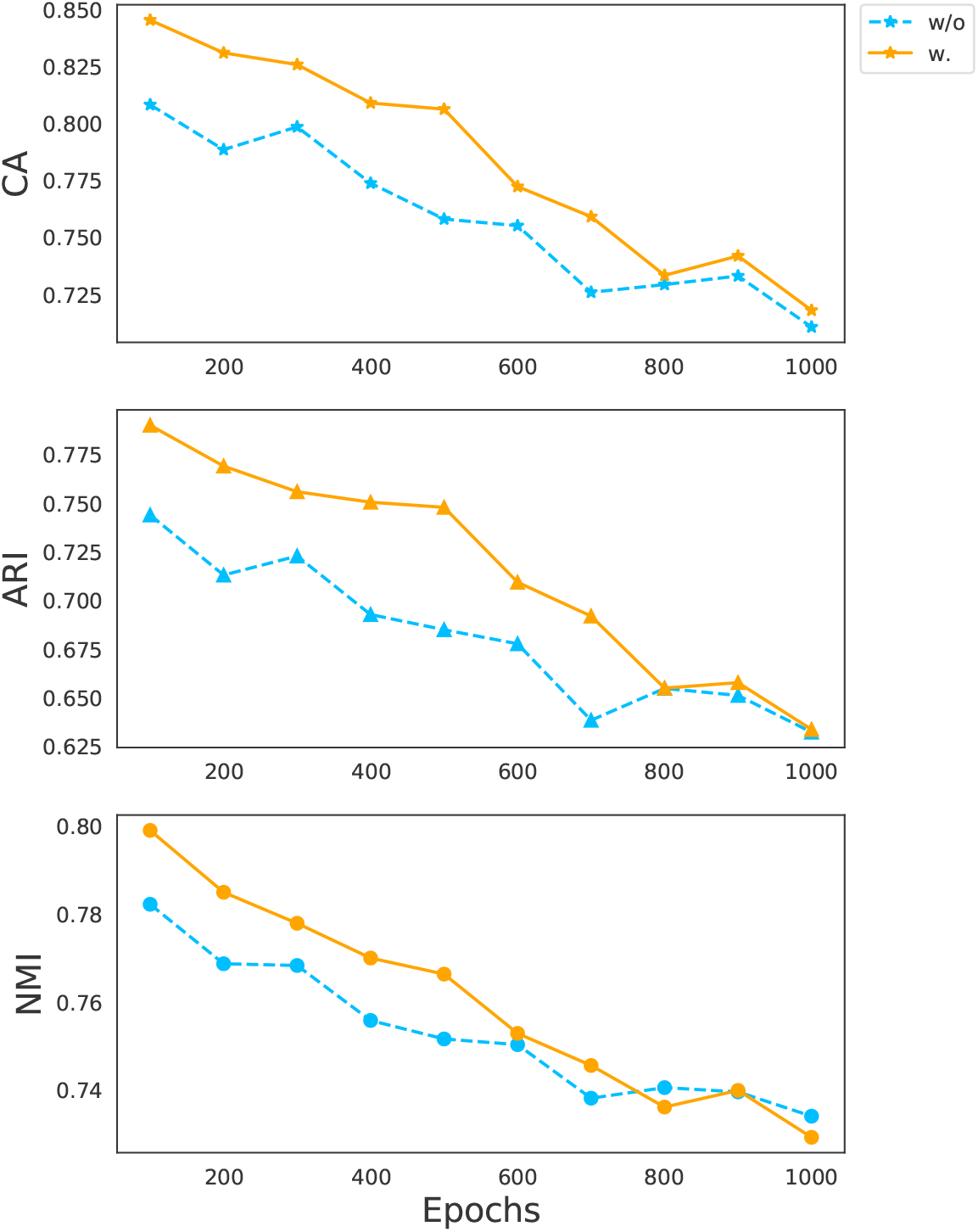
The comparison of clustering performances using deep clustering loss with or without consistency constraint.

With the training iteration going on, our proposed model can obtain better features, but this advantage will gradually weaken. This is mainly due to that the reconstruction task and the clustering task are under the different hypothesis, whose process is unsupervised and without real label information, and clustering loss can only optimize the joint distribution of latent features and it is difficult to divide the inaccurate potential.

## 5 Conclusions

Clustering is a fundamental analysis in scRNA-seq data analysis. In this paper, we introduced scDeepC3, a novel deep clustering model containing an AutoEncoder with adaptive shortcut connection and using deep clustering loss with consistency constraint for clustering analysis of scRNA-seq data. Empirical experiments demonstrate that scDeepC3 can effective extract embedded representations, which is optimized for clustering, of the high-dimensional input through a nonlinear mapping. The comparison of scDeepC3 with other clustering methods showcased that it always achieved better performance. In addition, the ablation studies intuitively demonstrate how scDeepC3 effectively improves the clustering results. From the experimental results, we can conclude that both adaptive shortcut connection layer and consistency constraint are useful to extract better embedded features for clustering. The results evaluated every 100 epochs also demonstrate that our method scDeepC3 has better and more stable training process.

In summary, scDeepC3 has shown to be a powerful tool for scRNA-seq clustering analysis. It can effective extract clustering-friendly embedded features. With the increasing popularity of scRNA-seq in biomedical research, we expect scDeepC3 will make better.

## Notes

### Competing Interest Statement

The authors have declared no competing interest.

## References

[1] C. Ziegenhain, B. Vieth, S. Parekh, B. Reinius, A. Guillaumet-Adkins, M. Smets, H. Leonhardt, H. Heyn Hellmann, and W. Enard, “Comparative analysis of single-cell rna sequencing methods,” Molecular cell, vol. 65, no. 4, pp. 631–643, 2017.

[2] W. Stephenson, L. T. Donlin, A. Butler, C. Rozo, B. Bracken, A. Rashidfarrokhi, S. M. Goodman, L. B. Ivashkiv, V. P. Bykerk, D. E. Orange et al., “Single-cell rna-seq of rheumatoid arthritis synovial tissue using low-cost microfluidic instrumentation,” Nature communications, vol. 9, no. 1, pp. 1–10, 2018.

[3] D. Talwar, A. Mongia, D. Sengupta, and A. Majumdar, “Autoimpute: Autoencoder based imputation of single-cell rna-seq data,” Scientific reports, vol. 8, no. 1, pp. 1–11, 2018.

[4] V. Y. Kiselev, T. S. Andrews, and M. Hemberg, “Challenges in unsupervised clustering of single-cell rna-seq data,” Nature Reviews Genetics, vol. 20, no. 5, pp. 273–282, 2019.

[5] J. Peng, X. Wang, and X. Shang, “Combining gene ontology with deep neural networks to enhance the clustering of single cell rna-seq data,” BMC bioinformatics, vol. 20, no. 8, p. 284, 2019.

[6] S. Wan, J. Kim, and K. J. Won, “Sharp: hyperfast and accurate processing of single-cell rna-seq data via ensemble random projection,” Genome Research, vol. 30, no. 2, pp. 205–213, 2020.

[7] J. A. Hartigan and M. A. Wong, “Algorithm as 136: A k-means clustering algorithm,” Journal of the royal statistical society. series c (applied statistics), vol. 28, no. 1, pp. 100–108, 1979.

[8] B. Wang, J. Zhu, E. Pierson, D. Ramazzotti, and S. Batzoglou, “Visualization and analysis of single-cell rna-seq data by kernel-based similarity learning,” Nature methods, vol. 14, no. 4, pp. 414–416, 2017.

[9] S. Park and H. Zhao, “Spectral clustering based on learning similarity matrix,” Bioinformatics, vol. 34, no. 12, pp. 2069–2076, 2018.

[10] R. Qi, J. Wu, F. Guo, L. Xu, and Q. Zou, “A spectral clustering with self-weighted multiple kernel learning method for single-cell rna-seq data,” Briefings in Bioinformatics, 2020.

[11] L. Parsons, E. Haque, and H. Liu, “Subspace clustering for high dimensional data: a review,” Acm Sigkdd Explorations Newsletter, vol. 6, no. 1, pp. 90–105, 2004.

[12] R. Zheng, Z. Liang, X. Chen, Y. Tian, C. Cao, and M. Li, “An adaptive sparse subspace clustering for cell type identification,” Frontiers in Genetics, vol. 11, p. 407, 2020.

[13] P. Indyk and R. Motwani, “Approximate nearest neighbors: towards removing the curse of dimensionality,” in Proceedings of the thirtieth annual ACM symposium on Theory of computing, 1998, pp. 604–613.

[14] C. Xu and Z. Su, “Identification of cell types from single-cell transcriptomes using a novel clustering method,” Bioinformatics, vol. 31, no. 12, pp. 1974–1980, 2015.

[15] V. Y. Kiselev, K. Kirschner, M. T. Schaub, T. Andrews, A. Yiu, T. Chandra, K. N. Natarajan, W. Reik, M. Barahona, A. R. Green et al., “Sc3: consensus clustering of single-cell rna-seq data,” Nature methods, vol. 14, no. 5, pp. 483–486, 2017.

[16] S. Park and H. Zhao, “Spectral clustering based on learning similarity matrix,” Bioinformatics, vol. 34, no. 12, pp. 2069–2076, 2018.

[17] L. v. d. Maaten and G. Hinton, “Visualizing data using t-sne,” Journal of machine learning research, vol. 9, o. Nov, pp. 2579–2605, 2008.

[18] E. Pierson and C. Yau, “Zifa: Dimensionality reduction for zero-inflated single-cell gene expression analysis,” Genome biology, vol. 16, no. 1, pp. 1–10, 2015.

[19] P. Lin, M. Troup, and J. W. Ho, “Cidr: Ultrafast and accurate clustering through imputation for single-cell rna-seq data,” Genome biology, vol. 18, no. 1, p. 59, 2017.

[20] W. V. Li and J. J. Li, “An accurate and robust imputation method scimpute for single-cell rna-seq data,” Nature communications, vol. 9, no. 1, pp. 1–9, 2018.

[21] C. Arisdakessian, O. Poirion, B. Yunits, X. Zhu, and L. X. Garmire, “Deepimpute: an accurate, fast, and scalable deep neural network method to impute single-cell rna-seq data,” Genome biology, vol. 20, no. 1, pp. 1–14, 2019.

[22] Y. LeCun, Y. Bengio, and G. Hinton, “Deep learning,” nature, vol. 521, no. 7553, pp. 436–444, 2015.

[23] G. Eraslan, L. M. Simon, M. Mircea, N. S. Mueller, and F. J. Theis, “Single-cell rna-seq denoising using a deep count autoencoder,” Nature communications, vol. 10, no. 1, pp. 1–14, 2019.

[24] Tian, J. Wan, Q. Song, and Z. Wei, “Clustering single-cell rna-seq data with a model-based deep learning approach,” Nature Machine Intelligence, vol. 1, no. 4, pp. 191–198, 2019.

[25] X. Li, K. Wang, Y. Lyu, H. Pan, J. Zhang, D. Stambolian, K. Susztak, M. P. Reilly, G. Hu, and M. Li, “Deep learning enables accurate clustering with batch effect removal in single-cell rna-seq analysis,” Nature communications, vol. 11, no. 1, pp. 1–14, 2020.

[26] J. Xie, R. Girshick, and A. Farhadi, “Unsupervised deep embedding for clustering analysis,” in International conference on machine learning, 2016, pp. 478–487.

[27] G. E. Hinton and R. R. Salakhutdinov, “Reducing the dimensionality of data with neural networks,” science, vol. 313, no. 5786, pp. 504–507, 2006.

[28] B. Wang, J. Zhu, E. Pierson, D. Ramazzotti, and S. Batzoglou, “Visualization and analysis of single-cell rna-seq data by kernel-based similarity learning,” Nature methods, vol. 14, no. 4, pp. 414–416, 2017.

[29] R. Lopez, J. Regier, M. B. Cole, M. I. Jordan, and N. Yosef, “Deep generative modeling for single-cell transcriptomics,” Nature methods, vol. 15, no. 12, pp. 1053–1058, 2018.

[30] C. H. Grønbech, M. F. Vording, P. N. Timshel, C. K. Sønderby, T. H. Pers, and O. Winther, “scvae: Variational auto-encoders for single-cell gene expression datas,” bioRxiv, p. 318295, 2018.

[31] D. P. Kingma and M. Welling, “Auto-encoding variational bayes,” arXiv preprint arXiv:1312.6114, 2013.

[32] P. Baldi, “Autoencoders, unsupervised learning, and deep architectures,” in Proceedings of ICML workshop on unsupervised and transfer learning, 2012, pp. 37–49.

[33] X. Lu, Y. Tsao, S. Matsuda, and C. Hori, “Speech enhancement based on deep denoising autoencoder.” In Interspeech, vol. 2013, 2013, pp. 436–440.

[34] K. He, X. Zhang, S. Ren, and J. Sun, “Deep residual learning for image recognition,” in Proceedings of the IEEE conference on computer vision and pattern recognition, 2016, pp. 770–778.

[35] O. Ronneberger, P. Fischer, and T. Brox, “U-net: Convolutional networks for biomedical image segmentation,” in International Conference on Medical image computing and computer-assisted intervention. Springer, 2015, pp. 234–241.

[36] H. W. Kuhn, “The hungarian method for the assignment problem,” Naval research logistics quarterly, vol. 2, no. 1-2, pp. 83–97, 1955.

[37] M. Abadi, P. Barham, J. Chen, Z. Chen, A. Davis, J. Dean, M. Devin, S. Ghemawat, G. Irving, M. Isard et al., “Tensorflow: A system for large-scale machine learning,” in 12th {USENIX} symposium on operating systems design and implementation ({OSDI} 16), 2016, pp. 265–283.

[38] D. P. Kingma and J. Ba, “Adam: A method for stochastic optimization,” arXiv preprint arXiv:1412.6980, 2014.

[39] P. A. Estévez, M. Tesmer, C. A. Perez, and J. M. Zurada, “Normalized mutual information feature selection,” IEEE Transactions on neural networks, vol. 20, no. 2, pp. 189–201, 2009.

[40] J. M. Santos and M. Embrechts, “On the use of the adjusted rand index as a metric for evaluating supervised classification,” in International conference on artificial neural networks. Springer, 2009, pp. 175–184.

